# Detailed Colocalization Analysis of A- and B-type Nuclear Lamins: a Workflow Using Super-Resolution STED Microscopy and Deconvolution

**DOI:** 10.1101/2024.09.17.613415

**Authors:** Merel Stiekema, Owen N. Gibson, Rogier J.A. Veltrop, Frans C.S. Ramaekers, Jos L.V. Broers, Marc A.M.J. van Zandvoort

## Abstract

The inner nuclear membrane is covered by a filamentous network, the nuclear lamina, consisting of A- and B-type lamins as its major components. The A- and B-type lamins form independent but interacting and partially overlapping networks, as demonstrated by previous super-resolution studies. The nuclear lamina in fibroblast cultures derived from laminopathy patients shows an increased segregation of the A- and B-type lamin networks, which can be quantitatively expressed by the Pearson’s Correlation Coefficient (PCC). Blurring and noise (convolution), however, significantly affect the quality of microscopy images, which led us to optimize the deconvolution approach for Confocal Scanning Laser Microscopy (CSLM) and Stimulated Emission Depletion (STED) microscopy images. For that purpose, the differences in using a theoretical, experimental, or semi-experimental Point Spread Function (PSF), an important parameter for deconvolution, was evaluated for its use in deconvolution of CSLM and STED microscopy images of double immunolabeled healthy and laminopathy patient fibroblasts. The semi-experimental is a new PSF introduced in this study, which combines the theoretical and experimental PSF to solve issues that arise from noisy PSF recordings due to very small and thereby low intensity fluorescent beads. From these deconvoluted images, the colocalization of the lamin networks could not only be quantified at the level of the nucleus as a whole, but also at a subnuclear level. The latter was achieved by dividing the nucleus into multiple equal rectangles using a custom-made ImageJ macro in Fiji. In this detailed analysis, we found heterogeneity in the colocalization of lamins A/C and B1 within and between nuclei in both healthy and laminopathy dermal fibroblasts, which cannot be detected in one single analysis for the entire nucleus.

## 1. Introduction

A- and B-type lamins are the major components of the nuclear lamina, a filamentous network located between the inner nuclear membrane and (hetero-)chromatin (1,2). The nuclear lamina plays a fundamental role in the organization of the nuclear architecture in most mammalian cells (3,4). The major isoforms of the A-type lamins are lamins A and C, encoded by the *LMNA* gene (5). B-type lamins are composed of lamin B1 and lamin B2, encoded by *LMNB1* and *LMNB2*, respectively (6,7). Mutations in these genes, particularly the *LMNA* gene, cause a number of rare diseases collectively called laminopathies (8,9). This is a heterogeneous group and comprise, amongst others, cardiomyopathies, lipodystrophy, and the Hutchinson-Gilford progeria syndrome.

In healthy cells, the A- and B-type lamins form independent but interacting and overlapping networks, as demonstrated by previous studies using super-resolution microscopy (10–12). Additionally, super-resolution microscopy revealed that lamin B1 is located closer to the inner nuclear membrane, while lamins A/C are localized closer to the nucleoplasm (11,13). Cells obtained from laminopathy patients show alterations in the localization of each lamin subtype in the nuclear lamina, resulting in readily visible abnormalities, such as nuclear blebs, honeycomb-like structures, and donut-shaped nuclei (14). However, only a minor fraction of diseased cells shows these abnormal nuclear morphologies (15). In our previous study we revealed a significant decrease in the degree of colocalization of A- and B-type lamins when comparing dermal fibroblasts from laminopathy patients with control cells (16). This study also demonstrated the suitability of the indirect immunofluorescence staining approach in combination with Stimulated Emission Depletion (STED) microscopy for studying the different lamin subtypes in laminopathy cells, thereby circumventing the need for more labour intensive and artifact sensitive procedures, such as transfection protocols. We concluded that STED super-resolution light microscopy in combination with immunofluorescence staining protocols provides an excellent tool for further research into the lamin organization in healthy and diseased cells.

However, microscopy images are not exact copies of the object under the microscope, since the object image is affected by blurring and noise (17). This is explained by convolution, which means that the image arises from the convolution of the object and the Point Spread Function (PSF). The PSF describes how the microscope, determined by stable factors from the optics of the imaging system, but also by variable factors such as laser power and fluorescent behaviour of the sample, projects a small fluorescent point into the image plane (18,19). When the PSF is known, deconvolution of the image can take place, resulting into a better approximation of the original image with improved resolution and contrast (20). Additionally, the PSF is a measure for the quality of an optical system. The PSF can be theoretically modelled and calculated according to the properties of the optics involved, such as lens Numerical Aperture (NA), the emission wavelength of used fluorophores, and pinhole size. From now on we call this the theoretical PSF. Another option is determining an experimental PSF, which is often done by acquiring z-stacks from beads of known size (21). These must be sub-resolution beads, meaning that these should have a size below the theoretical resolution of the microscope. After acquiring a z-stack, the PSF can be distilled from the three-dimensional (3D) images with various software programmes. The shape of the PSF indicates the presence of potential deviations in the optical system. Furthermore, the Full Width at Half Maximum (FWHM) of the PSF can be determined as a measure for the resolution of the microscope images. Importantly, the distilled PSF can be used during the deconvolution process (17). As the experimental PSF, but not the theoretical PSF, takes deviations unique to the used microscope system into account, it brings the object restored from the microscope image closer to the true object. For this, the z-stacks of the sub-resolution beads should be made with microscope settings similar to those used for the actual imaging of the sample (22).

In the literature limited hands-on information is available on how to acquire an experimental PSF, although Cole *et al.* (19) published a thorough protocol for confocal scanning laser microscopy (CSLM), while additional information can also be found on commercial microscopy websites, such as that of Scientific Volume Imaging (22). Moreover, one of the working groups of the Quality Assessment and REProducibility for instruments and images in Light Microscopy (QUAREP-LiMi; https://quarep.org/) initiative, aims to define sample preparation, image acquisition, and data analysis protocols for testing resolution, including the use of sub-resolution fluorescent beads (23). Its focus is mainly on quality control and not the use in deconvolution. Despite these efforts, so far, no protocols for experimental PSF determination in STED super-resolution microscopy have been published.

While the determination of experimental PSF for confocal microscopy is rather straightforward, determining the experimental PSF for super-resolution microscopy is challenging. This is due to the very small size of the sub-resolution beads, which is inherently accompanied by low fluorescent intensities and high bleaching rates. In this study we introduce a semi-experimental PSF, which is modelled based on the microscopic parameters determined with use of the z-stack of sub-resolution beads, and is thus a combination of the theoretical and experimental PSF.

The aim of the underlying study was to optimize the deconvolution protocol for the colocalization study of A- and B-type lamins in healthy and diseased cells using the optimal PSF. For that purpose, we have evaluated the differences in outcome when using different PSFs (theoretical, experimental, or semi-experimental) in deconvolution of CSLM and STED microscopy images of fibroblasts immunofluorescently labeled for lamins A/C and B1. The best suitable PSF has been used for deconvolution. Subsequently, the colocalization of the lamins A/C and B1 networks was determined and expressed as the Pearson’s Correlation Coefficient (PCC). This is not only done at the level of the nucleus as a whole, but also at a subnuclear level, by dividing a nucleus into multiple rectangles that are separately analyzed, allowing for a more detailed analysis of the colocalization of lamins A/C and B1.

## 2. Materials and Methods

### 2.1 Cell culture

Normal human dermal fibroblasts (NHDF) were obtained from PromoCell (Heidelberg, Germany). Laminopathy patient dermal fibroblasts with an *LMNA* c.1130G>T (p.(Arg377Leu)) variant were obtained from a skin biopsy (see (16)) after written consent and permission from the Medical Ethical Committee of the Maastricht University Medical Centre (METC permission number 21-017). NHDF and laminopathy patient dermal fibroblasts were cultured in Dulbecco’s Modified Eagle Medium (DMEM; Biowest, Nuaillé, France) containing 10% fetal calf serum (FCS) (Gibco, Waltham, MA, USA) and 50 µg/ml Gentamycin (Dechra, Northwich, UK) at 37°C and 5% CO_2_ in a humified incubator. At confluency, the cells were trypsinized using 0.125% Trypsin (Gibco, Waltham, MA, USA)/0.02% EDTA/0.02% glucose solution in Phosphate Buffered Saline (PBS) and passaged at a 1:2 or 1:3 ratio. The fibroblasts were fixed and stained at passage number 5-9.

### 2.2 Cell fixation and immunofluorescence staining

Cells were seeded onto 18 mm round glass coverslips (#1.5; VWR, Radnor, USA) at a 1:2 or 1:3 ratio, grown for at least 48 hours (h), and fixed with 4% formaldehyde (Merck, Darmstadt, Germany) in PBS at room temperature (RT) for 15 min. The cells were stored at 4°C in PBS containing 0.01% sodium azide for a maximum of 7 days.

Prior to immunofluorescence (IF) staining, formaldehyde fixed cells were permeabilized in 0.1% Triton X-100 (AppliChem, Darmstadt, Germany) in PBS for 15 min at RT, followed by washing with PBS (3 x 3 min). Primary antibodies, diluted in PBS containing 3% Bovine Serum Albumin (BSA; Roche Diagnostics, Basel, Switzerland), were applied onto the coverslips and incubated for 1 h at RT. The following primary lamin antibodies were used: 1) Mouse monoclonal IgG1 anti-lamins A/C culture supernatant (Jol2; dilution 1:50; provided by Prof. dr. C. Hutchison, Durham University, UK) for immunostaining of lamins A and C; 2) Rabbit polyclonal IgG anti-lamin B1 (ab16048; 1 mg/mL; dilution 1:800; Abcam, Cambridge, UK) for immunostaining of lamin B1. After incubation with primary antibodies, the coverslips were washed again with PBS (3 x 3 min) and the secondary antibodies, diluted in PBS containing 3% BSA, were applied to the coverslips and incubated for 1 h at RT. The following secondary antibodies were used: 1) Goat anti-mouse IgG Abberior Star Green (1 mg/mL; dilution 1:500; Abberior Instruments, Göttingen, Germany) as secondary antibody for detection of lamins A and C; 2) Goat anti-rabbit IgG Abberior Star 512 (1 mg/mL; dilution 1:250; Abberior Instruments, Göttingen, Germany) for detection of lamin B1. A final washing step with PBS (3 x 3 min) was performed before the coverslips were mounted on a microscopy glass slide with Tris-Glycerol DABCO mounting medium (90% glycerol, 20 mM Tris-HCl pH 8.0, 2% 1,4-di-azo-bicyclo-2(2,2,2)-octane (Merck, Darmstadt, Germany)), and sealed with nail polish. The slides were stored at 4°C until imaging, maximally 7 days after immunofluorescence staining.

### 2.3 Confocal Scanning Laser Microscopy (CSLM) and Stimulated Emission Depletion (STED) microscopy

The immunostained cell samples were imaged using CSLM and 2D-STED microscopy (Leica TSC SP8 STED microscope with LAS X software (Leica, Wetzlar, Germany)). 2D-STED microscopy applies emission depletion only in the xy-plane and not in the z-plane. Throughout this study when mentioning STED microscopy, we refer to 2D-STED microscopy. Imaging was performed with an HC PL APO CS2 100x/1.40 oil lens. Image acquisition was performed in xyz mode, with gating of 0.2-7.0 ns, format of 1024 x 1024 pixels, speed of 400 Hz, and pixel size of 30 x 30 nm. The cells were imaged in photon counting mode, with a gain of 10%, and line accumulation of 6-8 for CSLM images and 16 for STED microscopy images. Abberior Star Green was excited with a white light laser (WLL) at 488 nm and its emission detected at 493-545 nm. Abberior Star 512 was excited at 521 nm and detected at 526-565 nm, respectively. Depletion in the xy-plane was done with a 592 nm STED laser, using 50% laser power. For all images, a small z-stack of three sections (step size 0.10 µm) was generated to fulfil the Nyquist criterion (i.e. the minimal sampling density needed to capture all information from the microscope into the image (24)), which is necessary for optimal image deconvolution.

### 2.4 Determination and characterization of the experimental Point Spread Function (PSF)

Coverslips were coated with 0.01% Poly-L-Lysine (Sigma Diagnostics, St. Louis, USA) by a 5-minute incubation at RT, followed by 1 h of drying at RT. For CSLM PSF samples, 175 ± 5 nm green-fluorescent microspheres (PS-Speck Microscope Point Source Kit; Molecular Probes, Eugene, USA) were diluted 1:1000 in MilliQ and sonicated for 2 minutes. For STED microscopy PSF samples, 50 nm green-fluorescent nanoparticles (polydispersity index <0.2) (DiagPoly^TM^ Fluorescent Polystyrene Nanoparticles Green 50 nm; CD Bioparticles, Shirley, New York, USA) were diluted 1:10^5^ in PBS and sonicated for 5 minutes. The diluted PSF beads were added to the coated coverslip and dried for 4 h up to overnight at RT, followed by mounting onto a microscopy glass slide as described for the cell lines.

Z-stacks of the beads were made by applying imaging settings used for the secondary antibody Abberior Star Green, for both the CSLM and STED microscopy PSF measurements. The z-stack (step size 0.1 µm) was started at the focal plane below the beads and ended above, to assure that the complete image of the bead was made. The experimental PSF was distilled from the bead images with the Huygens PSF Distiller Wizard in Huygens Professional version 19.10 (Scientific Volume Imaging, The Netherlands, http://svi.nl). With the distillation of the STED microscopy experimental PSF, microscopic parameters were software-estimated (numerical aperture (NA), excitation beam fill factor, STED saturation, and STED immunity factor) and a new PSF was generated. As this PSF was based on microscopic parameters of experimental bead measurements and not on regular images, we refer to it as the semi-experimental PSF.

For all PSF’s, the FWHM of the PSF (i.e., the width of the peak at the half of its maximum value) was determined with the PSF FWHM Estimator in the Huygens software, using a Lorentzian fit.

### 2.5 Full Width at Half Maximum (FWHM) determination of the lamin layer

To estimate the lamin layer thickness in deconvoluted images, the average FWHM of the lamina was determined at the mid-level of the nucleus, using Fiji (26). For the quantitative analysis, the layer thickness was determined at five positions in each cell. Intensity curves were plotted through lines drawn perpendicular to the lamina and fitting this with a Gaussian distribution. The thickness was determined at the same position for the CSLM and STED images of lamins A/C and B1 after deconvolution with different PSFs. The formula 2√2(ln2 σ), with σ as Gaussian width parameter, was used to calculate the FWHM (25). The interobserver variation was assessed in our earlier study (16).

The FWHM of the lamin layer could not be correctly assessed in images that were not deconvoluted, as their Gaussian fitting is systematically affected by high background levels, particularly in the STED images (Supplemental Figure 1).

### 2.6 Deconvolution and determination of the Pearson’s Correlation Coefficient (PCC)

All cellular images were deconvoluted with Huygens Professional, using the Classic Maximum Likelihood Estimation (CMLE) algorithm, a quality threshold of 0.05, and maximum iterations of 100. The used PSF is indicated in the results. The microscopic parameters extracted from the metadata of the images were used, except for some parameters for STED microscopy that were estimated with the sub-resolution bead measurements (i.e. NA: 1.26, excitation beam fill factor: 0.60, STED saturation factor 9.85, STED immunity factor: 14%). For CSLM images of the dye Abberior Star Green, a signal-to-noise ratio (SNR) of 8.8 was used, while for Abberior Star 512 we used SNR 7.3. For STED microscopy images these were was 10 and 7.1, respectively, which was calculated with a representative image (SNR =√((maximum value image histogram)/(lowest value image histogram)) ∗ 3).

Colocalization analysis of the deconvoluted images of the nucleus as a whole was performed using Huygens Professional’s Colocalization Wizard, with a Region of Interest (ROI) drawn around the cell, Costes method for background estimation, and PCC as colocalization coefficient. The PCC measures the degree of correlative variation of two channels; the higher the value, the more correlated the two channels are, with a perfect correlation for a value of +1, perfect anti-correlation for -1, and the absence of a relationship at 0 (26). The formula for PCC is as follows: 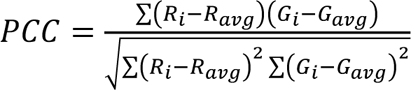 (26, 27). *R_avg_* and *G_avg_* refer to the mean intensities of the two channels to be analyzed (in this case detection of Abberior Star Green and Abberior Star 512), respectively. *R_i_* and *G_i_* refer to the intensity values of the Abberior Star Green and Abberior Star 512 channels of pixel *i*, respectively.

Since PCC values are not a linear scale, the average PCC of all cells within one cell line was calculated by first performing the Fisher *z*-transformation on the individual PCC values. The average of these *z*-values was calculated and back-transformed using the inverse Fisher transform to obtain the average PCC. The standard deviation (SD) of the *z*-values was added to (positive SD) or subtracted from (negative SD) the average *z*-value and subsequently back-transformed to a PCC value. The difference between those values and the average PCC were calculated to obtain the SD of the PCC.

Detailed subnuclear colocalization analysis using PCC was performed on each nucleus using a custom-made ImageJ macro in Fiji. First, a ROI of the outline of the nucleus (nuclear ROI) was made after using the threshold mean on the STED microscopy channel with emission detection of Abberior Star 512 (Figure 7A) (28). The CSLM and STED microscopy images were cropped around this ROI. The cropped image was divided into x^2^ rectangles (in which x varied between 4 and 12 to determine the optimal number of rectangles, but was 10 in the final analysis). The nuclear ROI and each rectangle were analyzed for the PCC (Rcoloc) with the colocalization threshold tool. Only rectangles with 100% overlap with the ROI of the nucleus were included in subsequent analysis.

For statistical analysis, the Mann-Whitney test was performed in GraphPad Prism (version 8.0.2). *p*-values are indicated in the text and *p*-values ≤ 0.05 were considered statistically significant.

## 3. Results

### 3.1 Deconvolution of CSLM and STED microscopy images

#### 3.1.1 Determination of the experimental and semi-experimental PSF

To perform deconvolution of the acquired CSLM and STED microscopy images of the nuclear lamina, the experimental PSF was acquired. This was distilled after averaging two 3D-images of multiple sub-resolution fluorescent beads (exact numbers in Table 1). The experimental PSF of CSLM does not demonstrate significant aberrations (e.g. spherical aberration) in the x-, y-, and z-plane (Figures 1A-C). The experimental STED microscopy PSF (Figures 1D-F) has a similar shape as compared to the CSLM PSF but is obviously much narrower in the x- and y-direction. However, around this spherically shaped PSF, a low intensity blur can be seen, mainly in the x, y- image (Figure 1D), most probably caused by background noise in the bead recording. When distilling the experimental PSF for STED microscopy, but not for CSLM, with the PSF Distiller Wizard of the Huygens software, some important microscopic parameters are estimated, specifically the NA, excitation beam fill factor, the STED saturation, and STED immunity factor. Based on the experimental PSF measurements and these subsequently extracted microscopic parameters, a new theoretical PSF can be generated, which we will refer to as the semi-experimental PSF (Figures 1G-I). Comparing this to the experimental PSF of STED microscopy, it demonstrates a similar spherical shape, but slightly more extended. Importantly, there is no blur surrounding this semi-experimental PSF and contains no noise because it is calculated.

**Table 1:** Lorentzian fitted Full Width at Half Maximum (FWHM) of the experimental and semi-experimental PSFs in the x-, y-, and z-plane. PSF of CSLM and STED microscopy is distilled after measurement of sub-resolution beads. For STED microscopy, the values for the experimental PSF and the semi-experimental PSF (i.e. theoretical PSF based on experimental microscopic parameters extracted from bead measurements) are indicated. For CSLM green-fluorescent microspheres (175 ± 5 nm) were measured, for STED microscopy green-fluorescent polystyrene nanoparticles (50 nm, polydispersity index <0.2) were used. The PSF is distilled after averaging multiple beads, collected from two separate measurements. CSLM experimental PSF used is visualized in Figures 1A-C, STED microscopy experimental in Figure D-F, STED microscopy semi-experimental in Figures 1G-I.

|  | CSLM experimental PSF | STED microscopy experimental PSF | STED microscopy semi-experimental PSF |
| --- | --- | --- | --- |
| **Number of beads** | 34 | 273 | 273 |
| **FWHM (nm) x-plane** | 181 | 82 | 95 |
| **FWHM (nm) y-plane** | 180 | 80 | 92 |
| **FWHM (nm) z-plane** | 397 | 405 | 588 |

**Figure 1:**
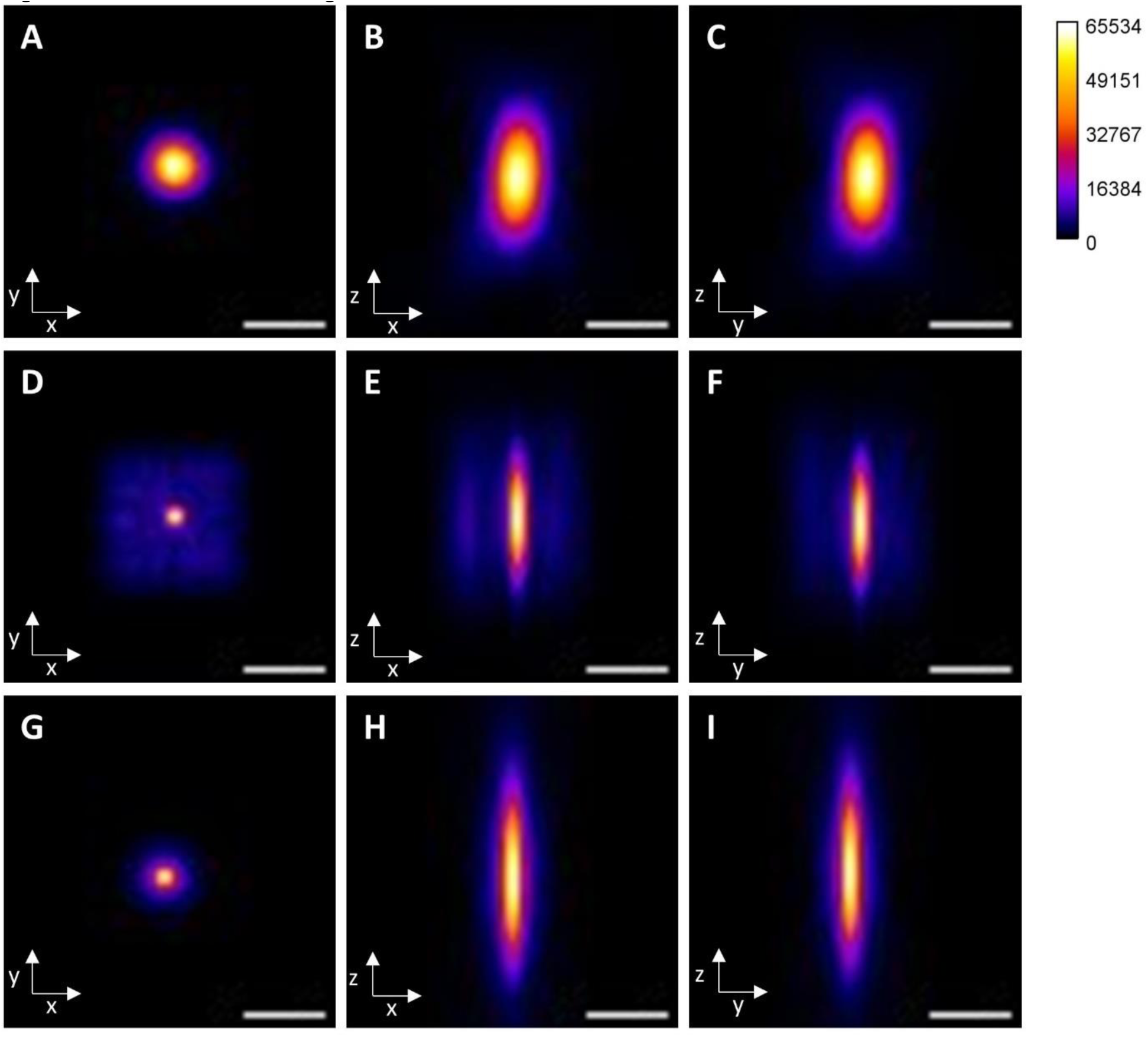
PSFs of CSLM and STED microscopy. **A-C**) Experimental PSF of CSLM, distilled form z-stacks of green- fluorescent microspheres (175 ± 5 nm), in the x,y- (**A**), x,z- (**B**), and y,z- (**C**) plane. **D-F**) Experimental PSF of STED microscopy, distilled from z-stacks of green-fluorescent polystyrene nanoparticles (50 nm, polydispersity index <0.2), in the x,y- (**D**), x,z- (**E**), and y,z- (**F**) plane. **G-I**) Semi-experimental PSF of STED microscopy (i.e. theoretical PSF based on experimental microscopic parameters extracted from bead measurements) in the x,y- (**G**), x,z- (**H**), and y,z- (**I**) plane. Scale bars: 0.5 µm. Calibration bar displays fluorescence intensities (scaled to maximum intensity value of 65534).

To determine the exact size of the PSF, the FWHM of intensity plots of different planes (x, y, and z) of the PSF was determined (Table 1). The FWHM of the STED microscopy PSFs is obviously much smaller in the x- and y-plane compared to the FWHM of CSLM, which is in line with the results seen in Figure 1. Since no 3D emission depletion was applied in the z-planes during STED image acquisition, the FWHM values of the CSLM and STED microscopy PSF are similar. The FWHM of the experimental PSF of STED microscopy is slightly smaller compared to that of the semi-experimental PSF, which is also visible as the smaller center of the PSF shapes in Figures 1D-F as compared to Figures 1G-I.

#### 3.1.2 Effect of different PSFs on deconvolution of CSLM images

The difference between performing no deconvolution, performing deconvolution with a theoretical PSF, or with an experimental PSF for CSLM images is displayed in Figures 2 and 3. These images show NHDF stained with antibodies against lamins A/C and lamin B1 (both visualized in (artificial) green) and imaged at the mid-level (Figure 2) or top (Figure 3) of the nucleus. The used theoretical PSF is computed by the software solely based on microscopic parameters extracted from the image and is thus not based on bead measurements, while the experimental PSF is based on bead measurements.

**Figure 2:**
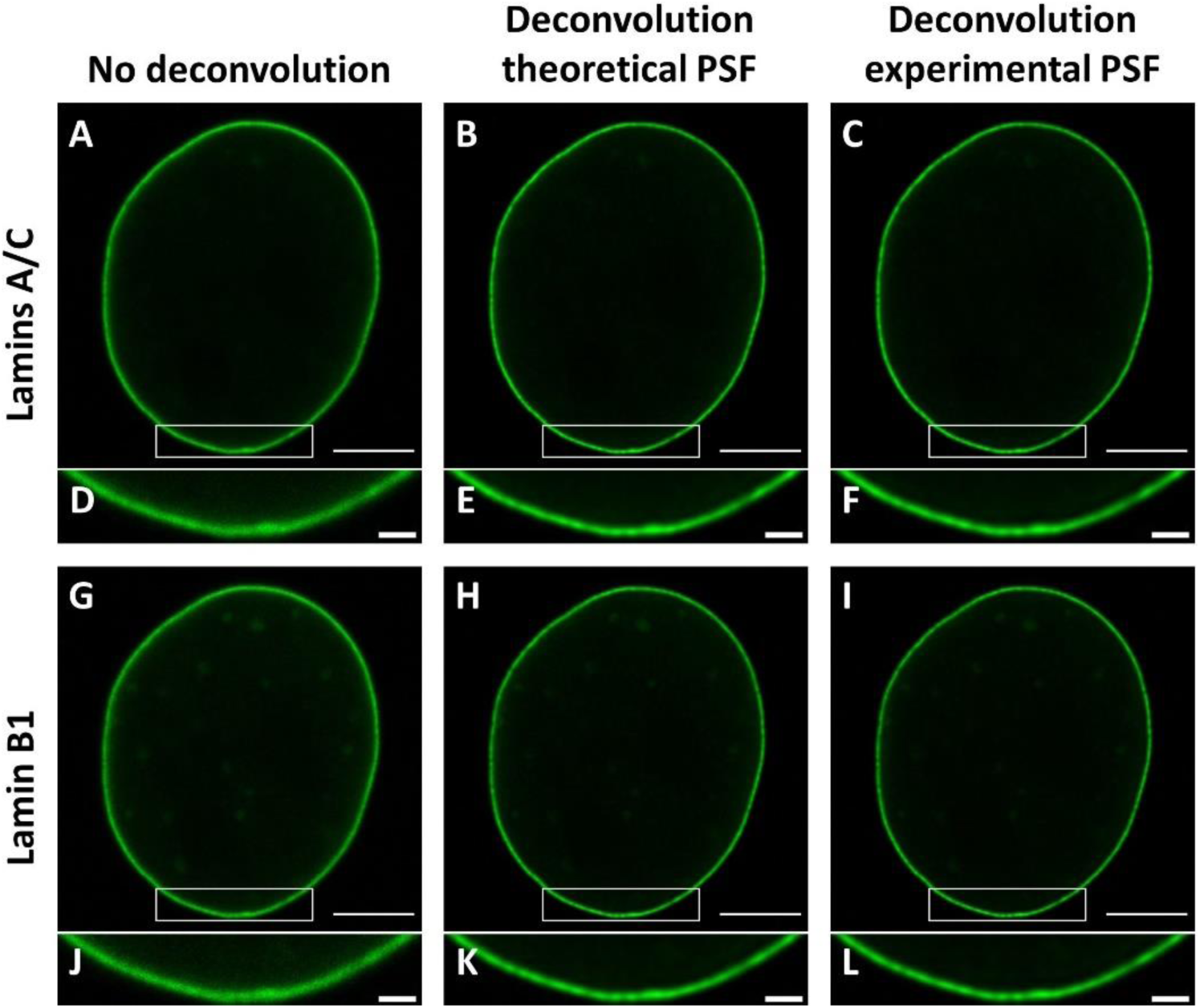
Effect of deconvolution using a theoretical or experimental PSF on CSLM images at the mid-level of the nucleus of normal human dermal fibroblasts (NHDF) stained with antibodies against lamins A/C (excitation at 488 nm, emission detection at 493-545 nm) and lamin B1 (excitation at 521 nm, emission detection at 526-565 nm). **A-C**) CSLM images of lamins A/C not deconvoluted (**A**), deconvoluted with a theoretical PSF (**B**), or with an experimental PSF (**C**). **D-F**) Higher magnification of the ROIs (white rectangles) in (**A-C**). **G-L**) CSLM images of lamin B1 not deconvoluted (**G**), deconvoluted with a theoretical PSF (**H**), or with an experimental PSF (**I**). **J-L**) Higher magnification of the ROIs in (**G-I**). Scale bars overview images: 5 µm. Scale bars higher magnification images: 1 µm.

**Figure 3:**
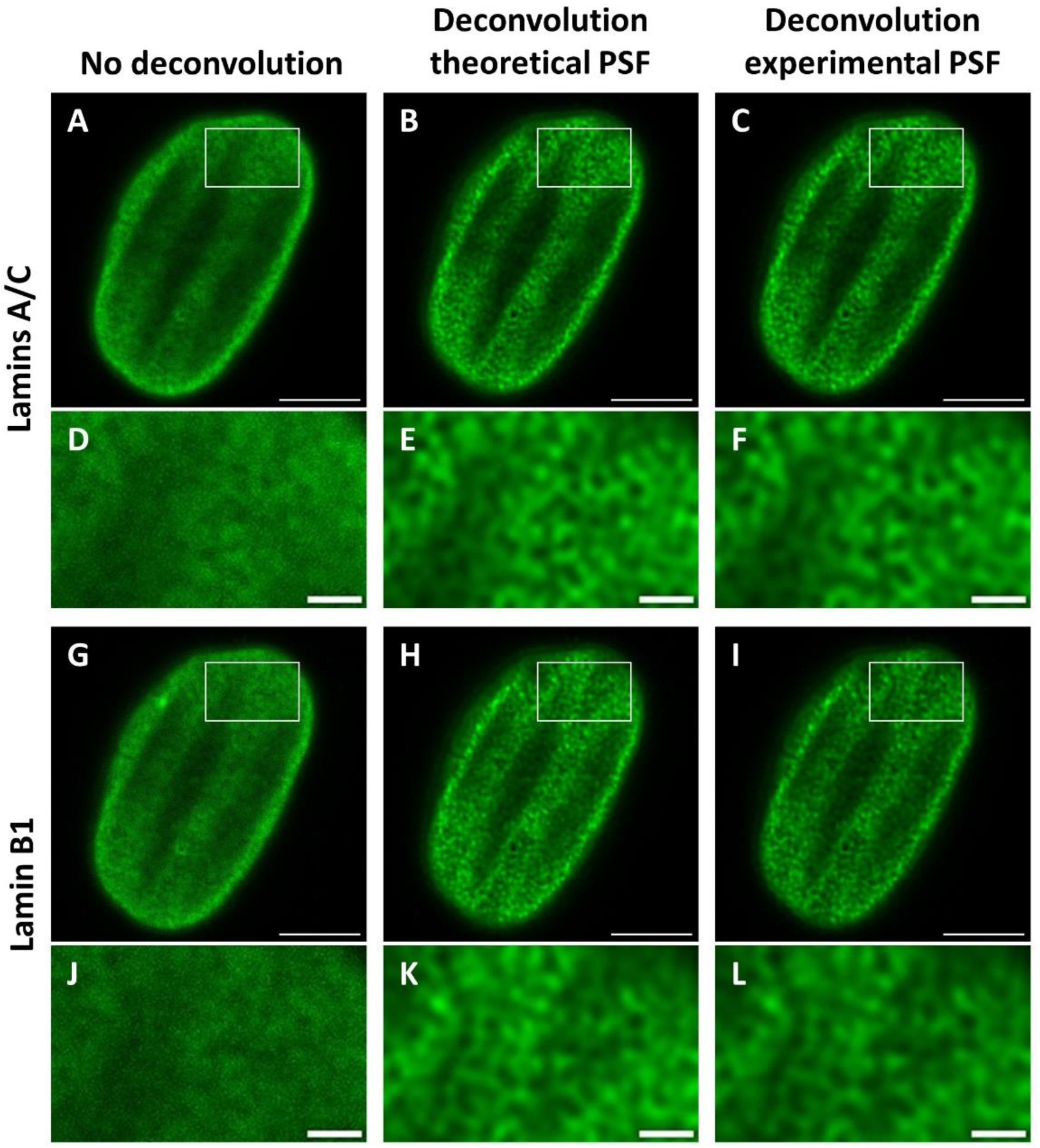
Effect of deconvolution using a theoretical or experimental PSF on CSLM images in the top of the nucleus of normal human dermal fibroblasts (NHDF) stained with antibodies against lamins A/C (excitation at 488 nm, emission detection at 493-545 nm) and lamin B1 (excitation at 521 nm, emission detection at 526-565 nm). **A-C**) CSLM images of lamins A/C not deconvoluted (**A**), deconvoluted with a theoretical PSF (**B**), or with an experimental PSF (**C**). **D-F**) Higher magnification of the ROIs (white rectangles) in (**A-C**). **G-L**) CSLM images of lamin B1 not deconvoluted (**G**), deconvoluted with a theoretical PSF (**H**), or with an experimental PSF (**I**). **J-L**) Higher magnification of the ROIs in (**G-I**). Scale bars images A-C and G-I: 5 µm. Scale bars higher magnification images D-F and J-L: 1 µm.

Comparing non-deconvoluted images to deconvoluted images of the mid-level of the nucleus by visual inspection (Figure 2), either deconvoluted with a theoretical or an experimental PSF, reveals that the lamin layer looks sharper and contains less noise after performing deconvolution. The top images of the nucleus (Figure 3) show a network that can be distinguished after performing deconvolution, compared to a vaguer image without deconvolution.

Measuring the lamina thickness of the deconvoluted images of the mid-level of the nucleus reveals a 5% thinner lamin layer in the images deconvoluted with the experimental PSF compared to those deconvoluted with the theoretical PSF (Supplemental Table 1). More importantly, the experimental PSF corrects for setup-specific physical deviations, so the experimental PSF is used for deconvolution of CSLM images in the rest of this study.

#### 3.1.3. Effect of different PSFs on deconvolution of STED microscopy images

The effect of applying deconvolution with different PSFs on STED microscopy images is visualized in Figures 4 and 5. The different PSFs that are compared include the theoretical PSF (based on microscopic parameters extracted from the image), the experimental PSF (distilled from sub-resolution bead measurements), and the semi-experimental PSF (based on experimental microscopic parameters extracted from sub-resolution bead measurements). Comparing non-deconvoluted to deconvoluted images, regardless of which PSF is used, demonstrates a sharper lamin layer in the deconvoluted images at the mid-level of the nucleus (Figure 4) and a more defined network at the top-level of the nucleus (Figure 5), as visible by eye.

**Figure 4:**
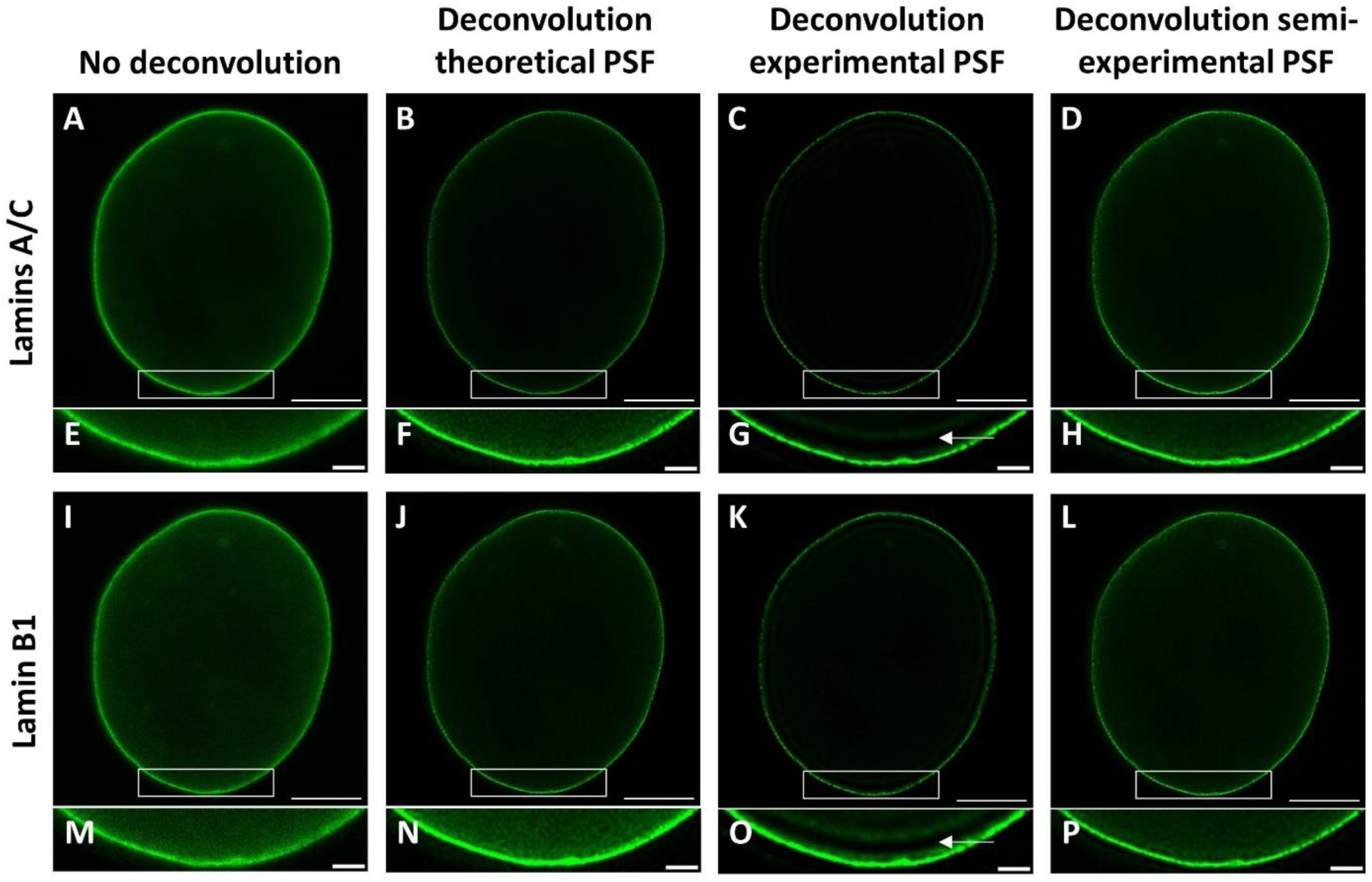
Effect of deconvolution using different PSFs on STED microscopy images at the mid-level of the nucleus of normal human dermal fibroblasts (NHDF) stained with antibodies against lamins A/C (excitation at 488 nm, emission detection at 493-545 nm) and lamin B1 (excitation at 521 nm, emission detection at 526-565 nm). **A-D**) STED microscopy images of lamins A/C not deconvoluted (**A**), deconvoluted with a theoretical PSF based (i.e. microscopic parameters extracted from the image) (**B**), deconvoluted with an experimental PSF distilled from bead measurements (**C**), or deconvoluted with a semi-experimental PSF (i.e. based on experimental microscopic parameters extracted from bead measurements) (**D**). **E-H**) Higher magnification and enhanced brightness of the ROIs (white rectangles) in (**A-D**). Arrow in (**G**) indicates ringing artefact of deconvolution. **I-L**) STED microscopy images of lamin B1 not deconvoluted (**I**), deconvoluted with a theoretical (**J**), deconvoluted with an experimental PSF (**K**), or deconvoluted with a semi-experimental PSF (**L**). (**M-P**) Higher magnification and enhanced brightness of ROIs in (**I-L**). Arrow in (**O**) indicates ringing artefact of deconvolution. Scale bars overview images: 5 µm. Scale bars higher magnification images: 1 µm.

**Figure 5:**
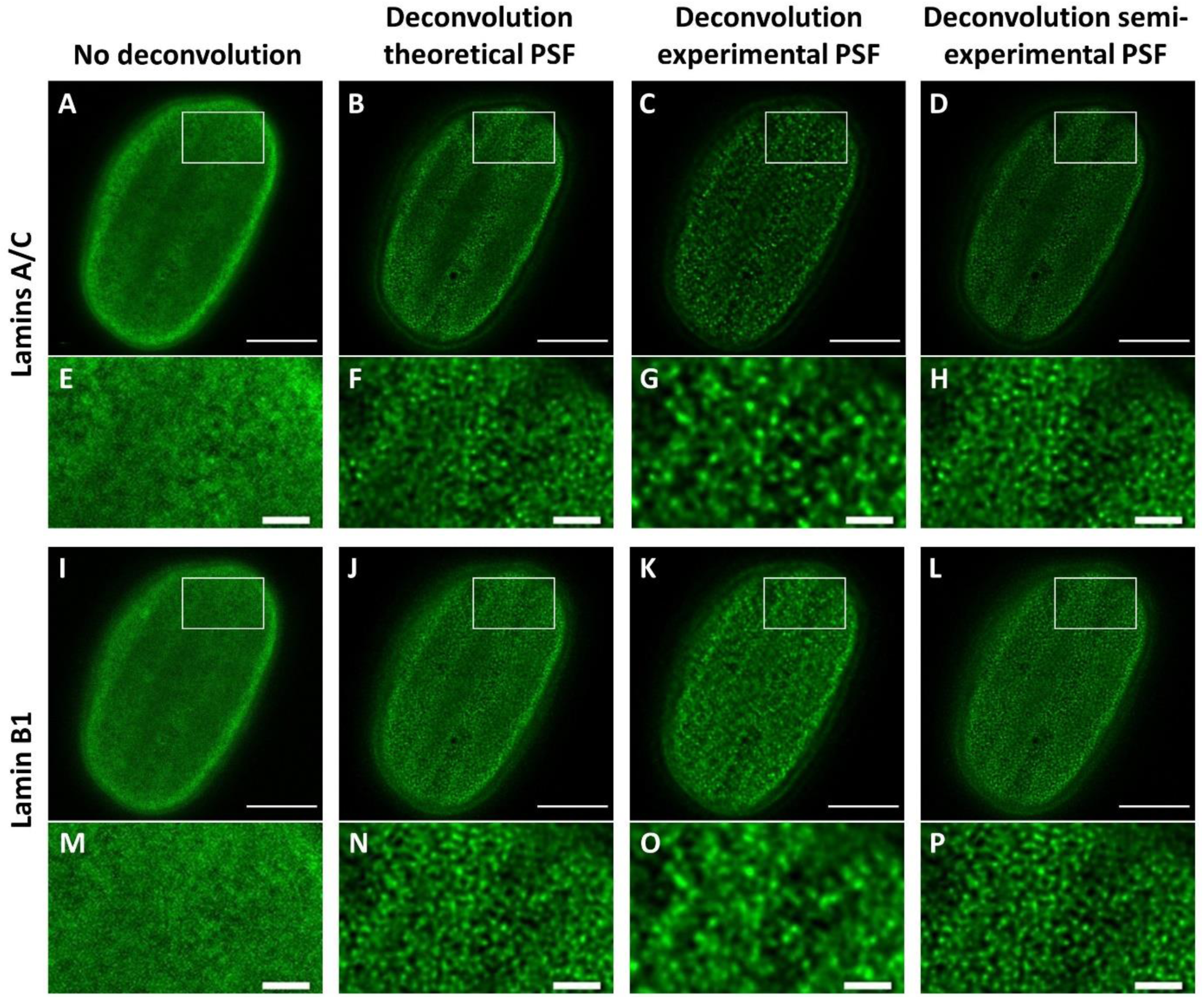
Effect of deconvolution using different point PSFs on STED microscopy images at the top of the nucleus of normal human dermal fibroblasts (NHDF) stained with antibodies against lamins A/C (excitation at 488 nm, emission detection at 493-545 nm) and lamin B1 (excitation at 521 nm, emission detection at 526-565 nm). **A-D**) STED microscopy images of lamins A/C not deconvoluted (**A**), deconvoluted with a theoretical PSF (i.e. based on microscopic parameters extracted from the image) (**B**), deconvoluted with an experimental PSF distilled from bead measurements (**C**), or deconvoluted with a semi-experimental PSF (i.e. based on experimental microscopic parameters extracted from bead measurements) (**D**). **E-H**) Higher magnification of the ROIs (white rectangles) in (**A-D**); (**G**) reveals over-emphasis on brighter structure and elimination of finer detailed structures. **I-L**) STED microscopy images of lamin B1 not deconvoluted (**I**), deconvoluted with a theoretical PSF (**J**), deconvoluted with an experimental PSF (**K**), or deconvoluted with semi-experimental PSF (**L**). (**M-P**) Higher magnification of ROIs in (**I-L**); (**O**) reveals over-emphasis on brighter structures and elimination of finer detailed structures. Scale bars overview images: 5 µm. Scale bars higher magnification images: 1 µm.

Deconvolution performed with different PSFs has a different visible outcome. Most strikingly, deconvolution performed with the experimental PSF reveals a ring with a lack of signal close to the lamin layer (arrows in Figures 4G and O). This is likely a so called ringing artifact, which has the appearance of dark and light ripples around bright features of an image and can occur by inadequate spatial sampling of the raw image or PSF, or a noisy image or PSF (29). As described above, the experimental PSF of STED microscopy contains a low intensity signal around the centre (Figure 1D). This artifact is not visible in the images deconvoluted with the theoretical PSF or the semi-experimental PSF.

At the top of the nucleus, deconvolution with the experimental PSF leads to over-emphasis of brighter structures and elimination of finer detailed structures, in contrast to the theoretical PSF or the semi-experimental PSF (Figure 5). Comparing deconvolution using the theoretical PSF and the semi-experimental PSF reveals, in this respect, minor to no differences amongst those two. The same comparison at the mid-level of the nucleus demonstrates a 25 - 27% thinner lamin layer in the images deconvoluted with the semi-experimental PSF compared to the theoretical PSF (Supplemental Table 1). Comparing the thickness of the lamin layer in images deconvoluted with the experimental PSF to the semi-experimental PSF reveals a similar thickness for images of lamins A/C, but a slightly (9%) thicker lamin layer in images of lamin B1 deconvoluted with the experimental PSF.

The use of the semi-experimental PSF in deconvolution thus leads to the best resolution. More important, the semi-experimental PSF is based on more setup-specific information, but does not lead to such obvious artefacts in the predicted structures and is thus considered the most reliable PSF. This semi-experimental PSF was used for the deconvolution of STED microscopy images in the rest of this study.

### 3.2 Detailed colocalization analysis of lamins A/C and B1

The images taken at the top of the fibroblast nuclei are used for analysis of the degree of colocalization of lamins A/C and lamin B1, since this view shows more of the lamin network than images taken at the mid-level of the nucleus (Figures 6A-F, lamins A/C visible in (artificial) red, lamin B1 in (artificial) green). In our previous study into the degree of colocalization of lamins A/C and B1 we compared the PCC of CSLM and STED microscopy images of the nucleus as a whole in NHDF and laminopathy patient dermal fibroblasts with an *LMNA* c.1130G>T (p.(Arg377Leu)) variant (16). In the current study, imaging settings of both CSLM and STED microscopy have been adapted (e.g. change to 100x/1.40 oil objective instead of 86x/1.20 objective) and the deconvolution method of the images was amended, to optimize the resolution. Therefore, both cell lines have been analyzed again for the degree of colocalization of lamins A/C and B1, now using the optimized imaging and deconvolution approaches as described above. To evaluate whether the higher resolution of STED microscopy indeed provides more information regarding the colocalization of lamins A/C and B1 as compared to images acquired with CSLM, we compared both systems. Figures 6G and H show that the observed degree of colocalization is lower in STED microscopy images as compared to CSLM images for both NHDF and the laminopathy patient fibroblasts. Importantly, only in the STED microscopy images the PCC of the laminopathy patient is significantly lower as compared to that of the NHDF. In CSLM images, the mean PCC is 0.92 (-0.04, +0.03) for the patient cells and 0.95 (-0.02, +0.02) for NHDF cells (*p*=0.08), while in STED microscopy images mean PCC values of 0.74 (-0.05, +0.04) and 0.83 (-0.04, +0.03) are found, respectively (*p*=0.002).

**Figure 6:**
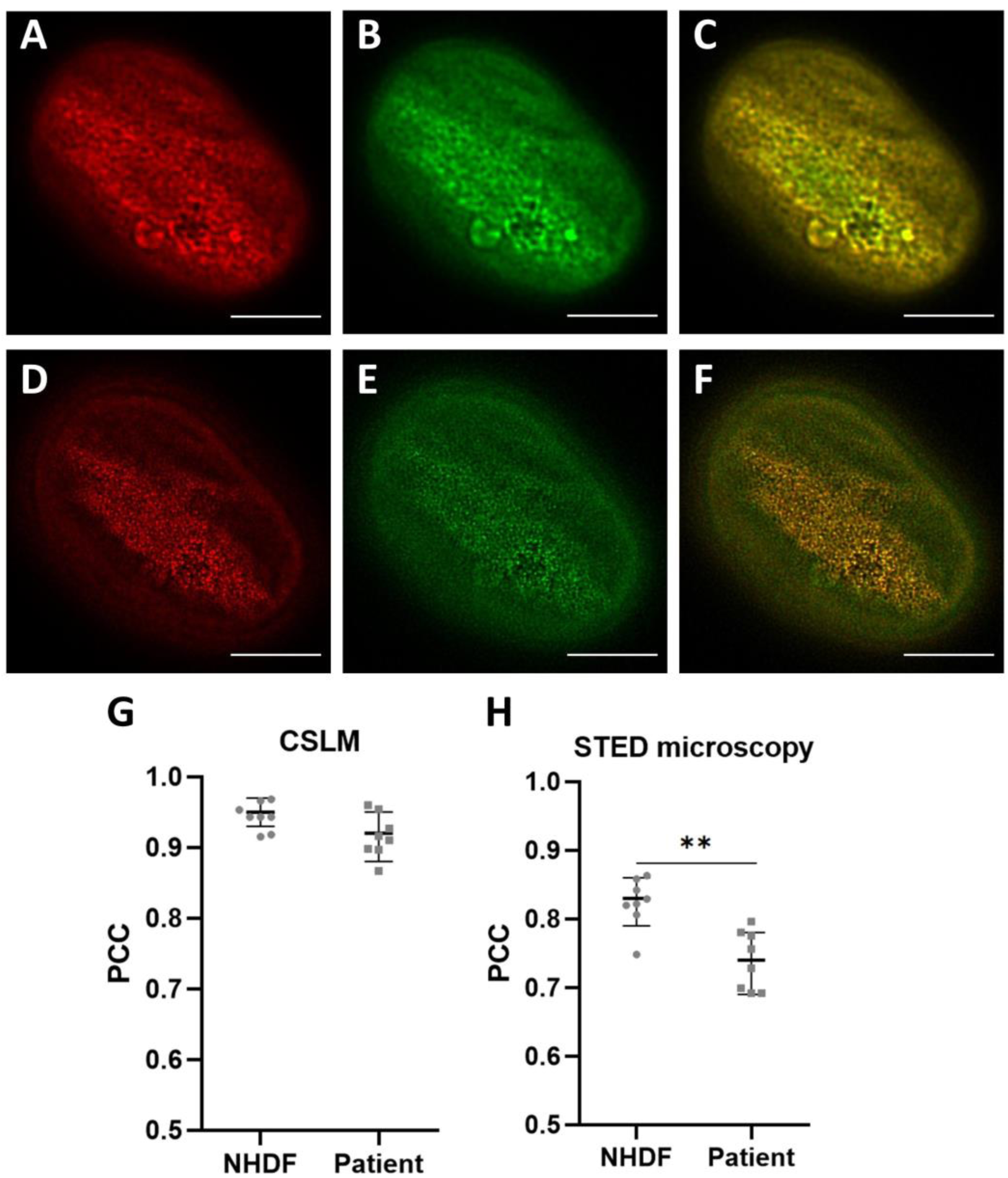
Comparison of the colocalization of lamins A/C and B1 in normal human dermal fibroblasts (NHDF) and the laminopathy patient cell line. **A-C**) CSLM images, deconvoluted with the experimental PSF, taken at the top of the nucleus of the NHDF; (**A**) lamins A/C in (artificial) red, (**B**) lamin B1 in (artificial) green, (**C**) merged image of (**A**) and (**B**). **D-E**) STED images, deconvoluted with the semi-experimental PSF, taken at the top of the NHDF nucleus; (**D**) lamins A/C in (artificial) red, (**E**) lamin B1 in (artificial) green, (**F**) merged image of (**D**) and (**E**). **G-H**) Degree of colocalization of lamins A/C and B1 in NHDF as compared to laminopathy patient dermal fibroblasts, expressed as the Pearson’s correlation coefficient (PCC); **G**) PCC of deconvoluted CSLM images using the experimental PSF. **H**) PCC of deconvoluted STED microscopy images using the semi-experimental PSF. The PCCs were determined using Huygens Professional. PCC values of each of the 8 nuclei analyzed per cell line are plotted separately with the mean ± SD indicated. **p≤ 0.01. Scale bars: 5 µm.

However, the colocalization analysis as described above (i.e. analysis of the top of the nucleus as a whole) does not provide information about more local, sub-nuclear heterogeneity in the degree of colocalization of lamins A/C and lamin B1. Therefore, a more detailed approach to analyze the top of the nuclei was applied, which could provide information about more localized changes of the lamina network. For this purpose, a symmetrical raster was placed over the images of the nuclei, dividing the images into multiple, equal regions (rectangles). For each area within such a rectangle the PCC of lamins A/C and B1 was determined (Figures 7B and C). In this analysis, only rectangles that were completely filled with a part of the nucleus (e.g. blue rectangle in Figure 7B) were included, while rectangles that did not or only partly cover an area of the nucleus were excluded (e.g. red rectangle in Figure 7B). First, we determined the number of rectangles that a nucleus should optimally be divided into, with 16, 25, 36, 49, 64, 81, 100 and 144 rectangles being used to overlay a nucleus (Figures 7D and E). The more rectangles are analyzed, the larger the spread in PCC values, but this reaches a plateau around 100 rectangles. Therefore, we chose to perform the detailed colocalization analysis of lamins A/C and B1 by dividing the nuclei into 100 rectangles.

**Figure 7:**
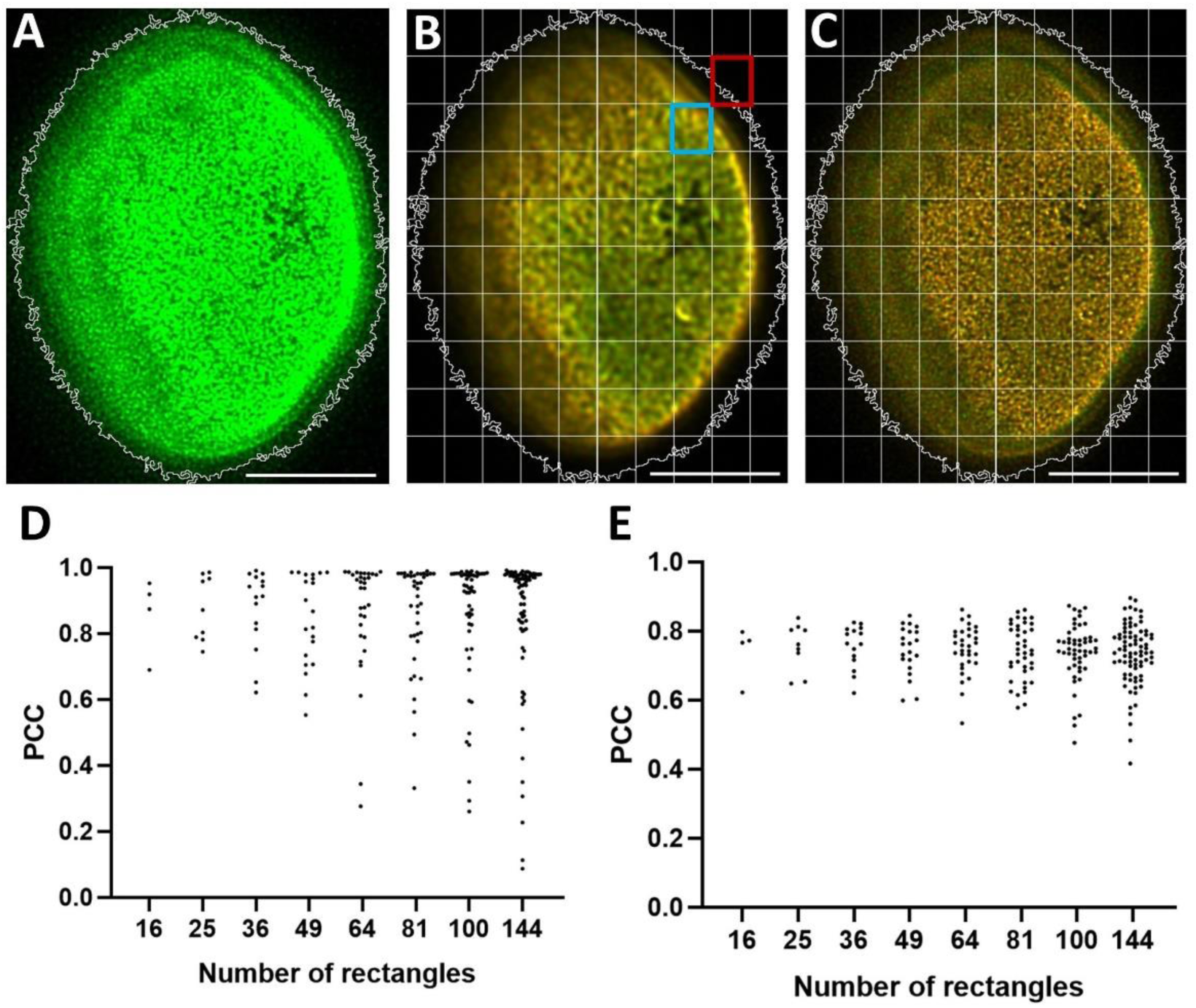
Detailed approach of colocalization analysis of lamins A/C and B1. **A**) STED microscopy image of lamin B1 with enhanced brightness to demonstrate the positioning of the ROI around the nucleus (winding white line) based on threshold mean in Fiji. **B**) CSLM image of top of the nucleus of NHDF with a ROI (winding white line) around the nucleus and 100 equal rectangles (straight white lines) that divide the image. Only rectangles that completely contain a part of the nucleus are included. Blue rectangle: example of rectangle that is included in the analysis. Red rectangle: example of rectangle that is excluded from the analysis. **C**) As A, but for a STED microscopy image; **D**) Different Pearson’s correlation coefficient (PCC) values of included rectangles when performing the analysis on 16, 25, 36, 49, 64, 81, 100 or 144 equal rectangles, of which 4, 6, 15, 21, 32, 40, 52, and 80 rectangles were included, respectively, in the CSLM image of (**B**). **E**) Similar as (**D**), but for the STED microscopy image of (**C**). Scale bars: 5 µm.

The detailed colocalization analysis of lamins A/C and B1 using 100 rectangles was performed for both CSLM and STED microscopy images of 8 NHDF and 8 laminopathy patient fibroblasts (Figure 8). Already at a first glance, it is obvious that there is a difference between NHDF and the laminopathy patient fibroblasts, as the majority of the local PCC values is much lower in the patient nuclei. Additionally, the CSLM images (Figures 8A and C) reveal higher PCC values compared to STED microscopy images (Figures 8B and D), confirming the ability of STED microscopy to better demonstrate differences in colocalization of lamins A/C and B1. Comparing the analysis of the nucleus as a whole to the detailed analysis, both determined in Fiji (Supplemental Figure 2), clearly demonstrates a significant local variation in the (absence of) lamins A/C and B1 colocalization in each nucleus, that is only visible in the detailed analysis. This is true for CSLM as well as STED microscopy images, but strikingly for both NHDF and laminopathy patient fibroblasts. Additionally, considerable differences in the degree of heterogeneity are visible between nuclei within the same cell line.

**Figure 8:**
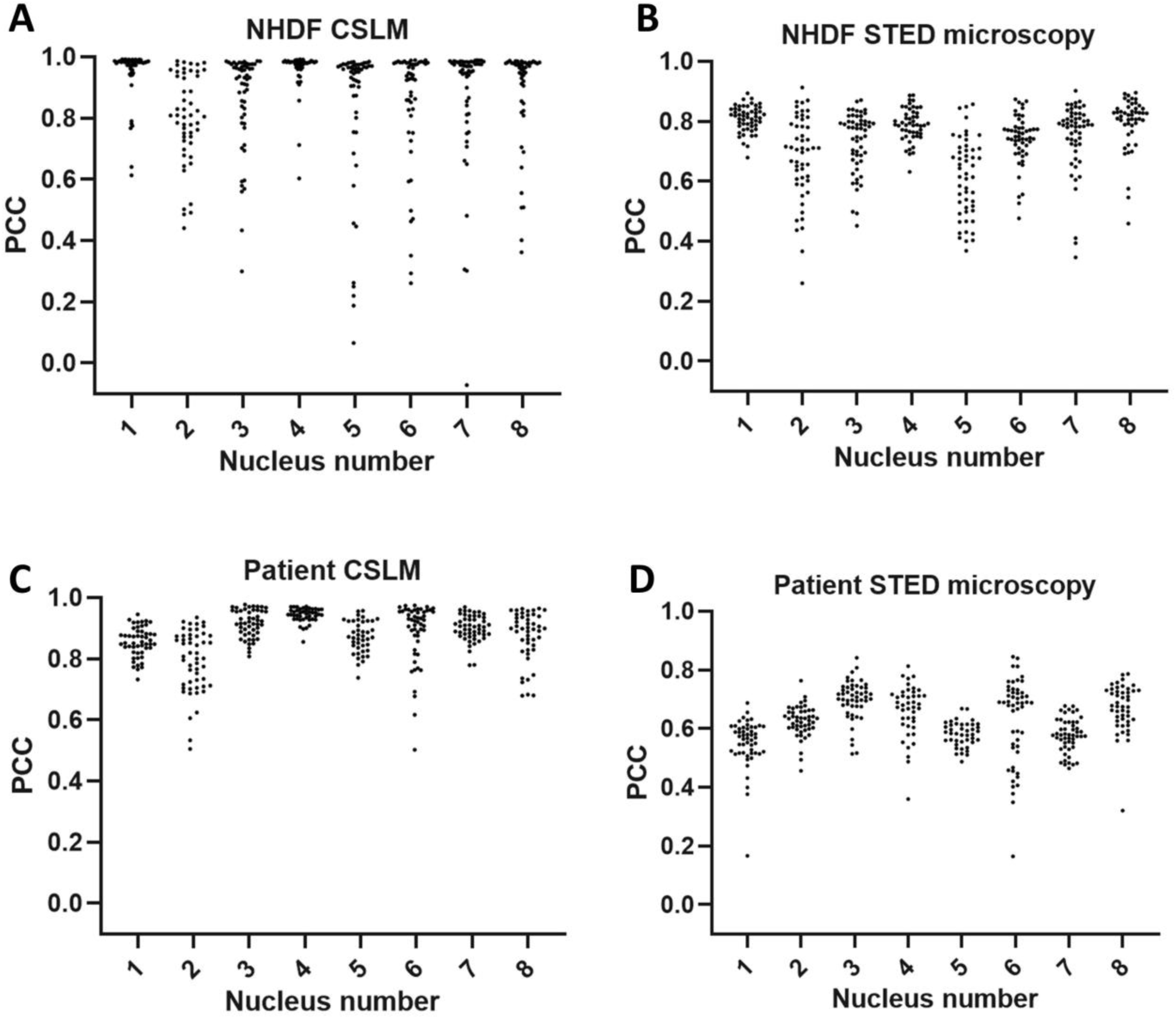
Colocalization of lamins A/C and B1 expressed as the detailed Pearson’s correlation coefficient (PCC). Microscopy images were divided into 100 equal rectangles and only rectangles with 100% overlap with the ROI of the nucleus were included in the colocalization analysis. (**A-B**) PCC of 8 CSLM (**A**) and 8 STED microscopy (**B**) images of NHDF. (**C-D**) PCC of 8 CSLM (**C**) and 8 STED microscopy (**D**) images of laminopathy patient dermal fibroblasts. X-axis of graph represents separate nuclei (images) numbered from 1-8. The PCC was determined using Fiji. Dots represent the individual PCCs of the included rectangles.

To visualize the heterogeneity of lamins A/C and B1 colocalization within individual nuclei, heatmaps were generated (Figure 9), projecting the PCC values included in the dot plots of Figure 8 into each rectangle. In general, heatmaps belonging to CSLM images display higher PCC values compared to those of STED microscopy images. The distribution of the PCC values within one nucleus, however, follows largely the same trend when comparing the CSLM and STED microscopy images of the same nucleus, with the understanding that the PCC values of the STED microscopy images are overall lower due to the higher resolution of these images.

**Figure 9:**
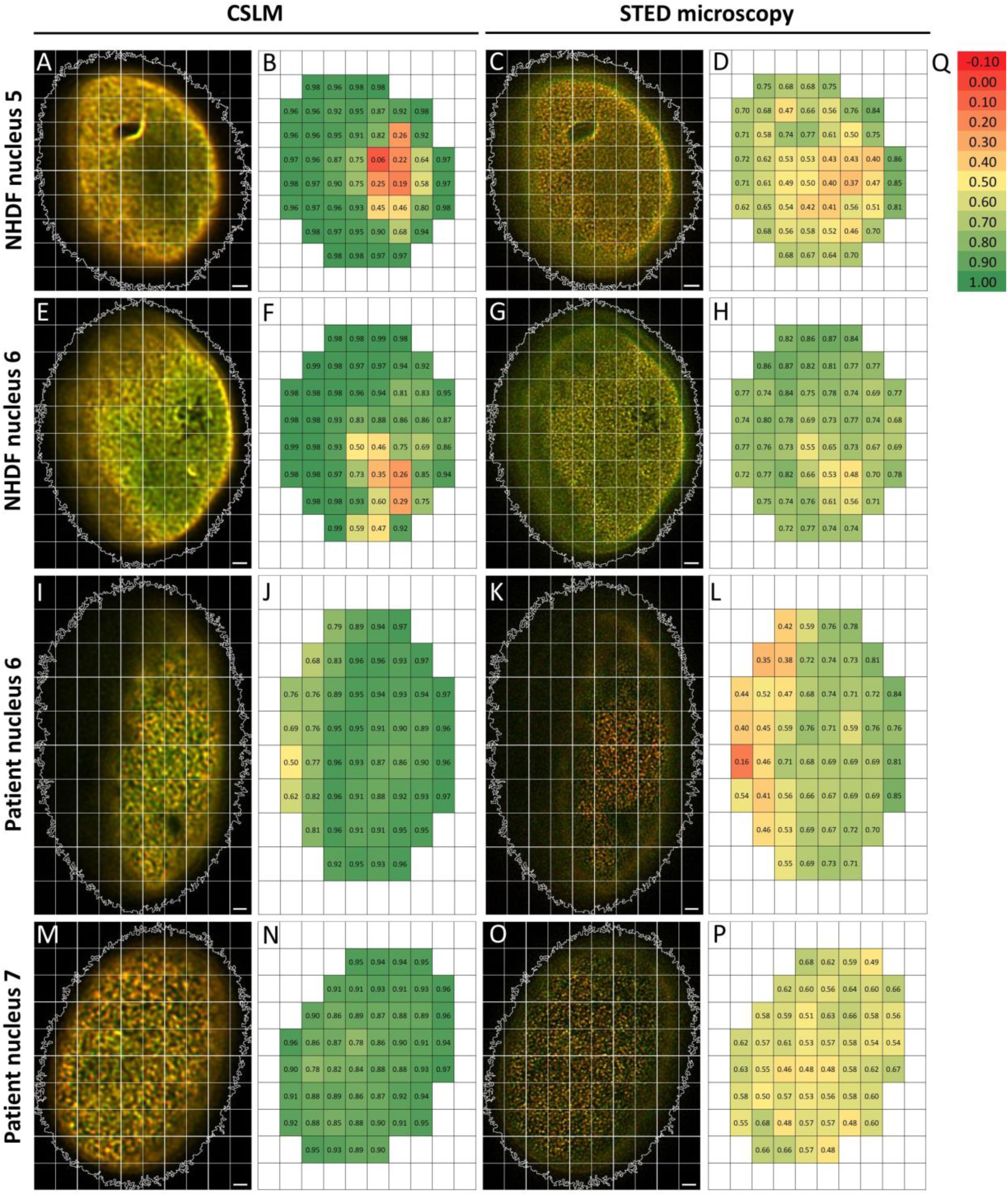
Heatmaps representing colocalization of lamins A/C and B1, expressed as the Pearson’s correlation coefficient (PCC) of NHDF (**A-H**) and of laminopathy patient dermal fibroblasts (**I-P**). (**A-D**) CSLM (**A-B**) and STED microscopy (**C-D**) image and related colocalization heatmap of NHDF nucleus number 5 represented in Figures 8 A and B. (**E-H**) CSLM (**E-F**) and STED microscopy (**G-H**) image and related colocalization heatmap of NHDF nucleus number 6 represented in Figures 8 A and B. (**I-L**) CSLM (**I-J**) and STED microscopy (**K-L**) image and related colocalization heatmap of the laminopathy patient nucleus number 6 represented in Figures 8 C and D. (**M-P**) CSLM (**M-N**) and STED microscopy (**O-P**) image and related colocalization heatmap of laminopathy patient nucleus number 7 represented in Figures 8 C and D. (**Q**) Color scale used for heatmaps. Rectangular ROIs in microscopy images represent the 100 equal rectangles that divide the image for the detailed colocalization analysis. The winding white line represents the outline of the nuclei, determined as shown in Figure 7. In the heatmaps only the PCC values of the rectangles with 100% overlap with the ROI of the nucleus are included. Scale bars represent 1 µm.

A more detailed inspection of the different NHDF nuclei analyzed in this heatmap approach shows that the heterogeneity of colocalization of lamins A/C and B1 differs considerably per imaged nucleus. Comparing NHDF nucleus number 5 and 6 for example, displays a broad variation in distribution of values in nucleus 5 (Figures 9A-D), but a more homogeneous distribution of PCC values in nucleus 6 (Figures 9E-H).

Also, for the laminopathy patient fibroblasts differences in the colocalization of lamins A/C and B1 are seen between different nuclei, as can be appreciated from Figure 8D. For instance, analysis of nucleus number 6 (Figures 9I-L) reveals a prominent variation in PCC values, while for nucleus number 7 (Figures 9M-P) this variation is less obvious. Examining those images in more detail (Figures 9I-P) reveals that most of the lower PCC values of number 6 (Figures 9I-L) are found on the left side of the nucleus. Although here the left side of the nucleus is less in focus compared to the right side, this does not explain the large variety in PCC values in this image. This becomes obvious when looking at the other images where some parts are less in focus, but PCC values do not vary. Finally, examining nucleus number 7 (Figures 9M-P) in more detail reveals that the regions with higher and lower lamins A/C and B1 colocalization are spread evenly across this nucleus.

We conclude that, while the detailed analysis of PCC values as shown in Figure 8 already provides valuable information on the variation of the colocalization of A-and B-type lamins over the nucleus, the heatmap approach adds significant morphological information by localizing this variation to specific nuclei and regions in the nucleus.

## 4. Discussion

In our previous study we demonstrated that STED microscopy in combination with indirect immunofluorescence protocols is an excellent tool for studying the nuclear lamina of laminopathy patient fibroblasts at high resolution (16). In this study, we further improved this approach by using an experimental or semi-experimental PSF for deconvolution, leading to an enhanced accuracy (20). We could furthermore obtain a more detailed and quantitative view of the distribution of the colocalization of lamins A/C and B1 by using sub-nuclear localized PCC values and heatmaps thereof.

### 4.1 (Semi-)experimental PSF for deconvolution of CSLM and STED microscopy images

The FWHM of the measured PSF provides information about the resolution of the microscope (19). For the CSLM settings the resolution in the x-direction was found to be 181 nm and 180 nm in the y-direction (Table 1). For STED microscopy this is 82 and 80 nm, respectively (experimental PSF). This decrease is as expected because the use of the STED depletion laser (30). The FWHM in the x- and y-direction of the STED experimental PSF is close to a previously reported STED lateral resolution between 30 and 80 nm for biological samples (31). The exact value is dependent on the properties of the sample and the applied depletion laser power. In the z-plane, there is no enhanced resolution for STED microscopy compared to CSLM, as the FWHM of CSLM is 397 nm, while this is 405 nm for STED microscopy. This is as expected with 2D-STED microscopy, as emission depletion is only applied in the x- and y-plane and not in the z-direction (32).

Deconvolution performed with an experimental PSF leads to a clear improvement of resolution and SNR for CSLM microscopy images (Figures 2 and 3). For STED microscopy images the same holds true, but in addition a ringing artifact is generated that can occur near sharp transitions in a signal (Figures 4 G and O) (29). A similar phenomenon can sometimes be seen in MRI images, here often referred to as truncation or Gibbs artifacts (33). This ringing artifact can be caused by the noise around the experimental PSF, as seen in Figure 1D.

The 50 nm beads necessary for sub-resolution bead measurements in STED microscopy suffer from low signal intensity and/or fast bleaching in acquiring z-stacks. This is more problematic compared to CSLM bead measurements (where larger beads are used), as the number of fluorescent molecules scales with the size of the beads (22). Therefore, there is a low SNR in the bead recordings with STED microscopy, which could lead to noise around the distilled experimental PSF. A comparison by Klemm *et al.* (21) demonstrated that gold beads have a much higher intensity and better SNR compared to fluorescent beads. However, using gold beads for PSF measurements requires imaging in reflection mode, which is not representative for the microscope settings when imaging a biological sample with fluorescent labeling. For optimal deconvolution, it is best to use microscopic imaging parameters that are as similar as possible to those used for the actual imaging of the sample (22). If PSF measurements are done solely to acquire information about the resolution of the optical set-up, gold beads can be preferred for a better SNR. However, if the PSF is also used for deconvolution, fluorescent beads are preferred.

In conclusion, to consider setup specific aberrations as much as possible, using the semi-experimental PSF determined using sub-resolution fluorescent beads was the preferred choice for deconvolution of STED microscopy images. Deconvolution performed with this semi-experimental PSF also leads to enhanced resolution and SNR in the STED microscopy images (Figures 4 and 5).

### 4.2 Heterogeneous colocalization of lamins A/C and B1

When analyzing the nucleus as a whole for the colocalization of lamins A/C and B1, expressed as the PCC, a significant (*p*=0.002) lower degree of colocalization is found in laminopathy patient dermal fibroblasts with an *LMNA* c.1130G>T (p.(Arg377Leu) variant as compared to NHDF, but only when using STED microscopy. In our previous study, we found a significant decrease in both CSLM and STED microscopy, but the difference was more apparent with STED microscopy (16), which indicates that the differences in the image acquisition and the deconvolution approach can affect the magnitude of the variation, but does not affect our basic conclusion. We confirmed again the ability of STED microscopy to demonstrate differences in the colocalization of lamins A/C and B1 better compared to CSLM. More importantly, the degree of lamins A/C and B1 colocalization was investigated in a more detailed manner in the current study. Analyzing multiple regions within an individual nucleus using sub-nuclear PCCs and heatmaps thereof, allowed the detection of regional differences in colocalization. In general, the degree of colocalization of lamins A/C and B1 is found to be heterogeneous within nuclei and between nuclei of both healthy and laminopathy patient fibroblasts. While part of these differences in laminopathy cells, such as herniations and honeycomb-like structures, already can be determined by visual inspection (15), this methods allows also to detect significant differences in colocalization in apparently normal subregions of these cells. The heterogeneity of lamins A/C and B1 colocalization is a novel finding that could not be detected when analyzing the nucleus as a whole, demonstrating the added value of this detailed colocalization analysis. Of course, this approach is not limited to this research topic and could also be combined with other software programs for image analysis and potentially be further automatized.

This heterogeneity in colocalization of lamins A/C and B1 within and between nuclei could potentially be related to differences in cell cycle, interaction with lamin-binding proteins, or chromatin interactions. This localized PCC analysis can thus be a very relevant addition to a broad range of studies. The heterogeneity of the colocalization of lamins A/C and B1 does support the current view that A- and B-type lamins form independent but interacting and overlapping networks (10–12). Additional healthy and diseased fibroblast cultures will have to be be analyzed to determine if a lower degree of colocalization of lamins A/C and B1 is a common phenotype in laminopathy patients. Moreover, by including patients with varying percentages of abnormal nuclei and different classifications of the *LMNA* mutation according to molecular genetics criteria (15), it can be assessed whether the degree of lamins A/C and B1 colocalization is correlated to the percentage of abnormal nuclei or the classification of the mutation.

In conclusion, the imaging and analysis workflow presented in this study allows for high-resolution and detailed colocalization analysis of lamins A/C and B1. The novel detailed colocalization approach provides valuable insights into the heterogeneity of the sub-nuclear colocalization of lamins A/C and B1 in both healthy dermal fibroblasts and in laminopathy patient fibroblasts. Follow-up studies will have to reveal if a lower degree of colocalization of A- and B-type lamins is a common phenotype in laminopathies and whether this information has potential clinical value. Finally, it may be obvious that the imaging and analysis workflow presented in this study not only allows for high-resolution and detailed colocalization analysis of lamins, but is applicable for association studies of cellular constituents and processes in a much broader sense.

## Data availability

The ImageJ script used for the data analysis is available upon request.

## Ethical approval

Laminopathy patient dermal fibroblasts were obtained and used for this study after written consent and permission from the Medical Ethical Committee of the Maastricht University Medical Centre (METC 21-017).

## Author contributions

Writing - original draft preparation: MS. Writing – review and editing: MS, OG, RJAV, FCSR, JLVB, MAMJvZ. Performing experiments: MS and OG. Supervision: FCSR, JLVB, and MAMJvZ. All authors contributed to the article and approved the submitted version.

## Acknowledgements

We thank dr. Johan A. Slotman (Department of Pathology, Optical Imaging Centre, Erasmus University Medical Centre, Rotterdam, The Netherlands) for valuable discussions concerning colocalization analysis and Ralph Hendriks for his help with the data analysis of the PCC. We acknowledge prof. dr. C. Hutchison (Durham University, UK) for providing the Jol2 anti-lamins A/C antibody. We acknowledge the support with STED microscopy by Helma Kuijpers and Kèvin Knoops from the Microscopy CORE Lab at the Faculty of Health, Medicine, and Life Sciences, Maastricht University.

