## Supplementary data for "Detailed Colocalization Analysis of A- and B-type Nuclear Lamins: a Workflow Using Super-Resolution STED Microscopy and Deconvolution"

***Supplemental Table 1:*** *Lamina thickness in CSLM and STED microscopy images of NHDF stained for lamins A/C and B1 and deconvoluted with different PSFs. The lamina thickness is determined by calculating the FHWM of intensity plots drawn perpendicular to the lamina as shown in Supplemental Figure 1. Values represent FWHM ± SD. The FWHM is determined at five positions in each cell, for 10 NHDF cells in total. Values were excluded if the height of the fitted curve is lower compared to the raw values (4 times in 500 determinations) (see example Supplemental Figure 1).*

| **PSF applied** | **CSLM lamins A/C** | **STED microscopy lamins A/C** | **CSLM lamin B1** | **STED microscopy lamin B1** |
|---|---|---|---|---|
| **Theoretical** | 241 ± 25 | 154 ± 38 | 246 ± 34 | 181 ± 38 |
| **Experimental** | 230 ± 22 | 115 ± 25 | 234 ± 35 | 144 ± 33 |
| **Semi-experimental** | N/A | 115 ± 20 | N/A | 132 ± 29 |

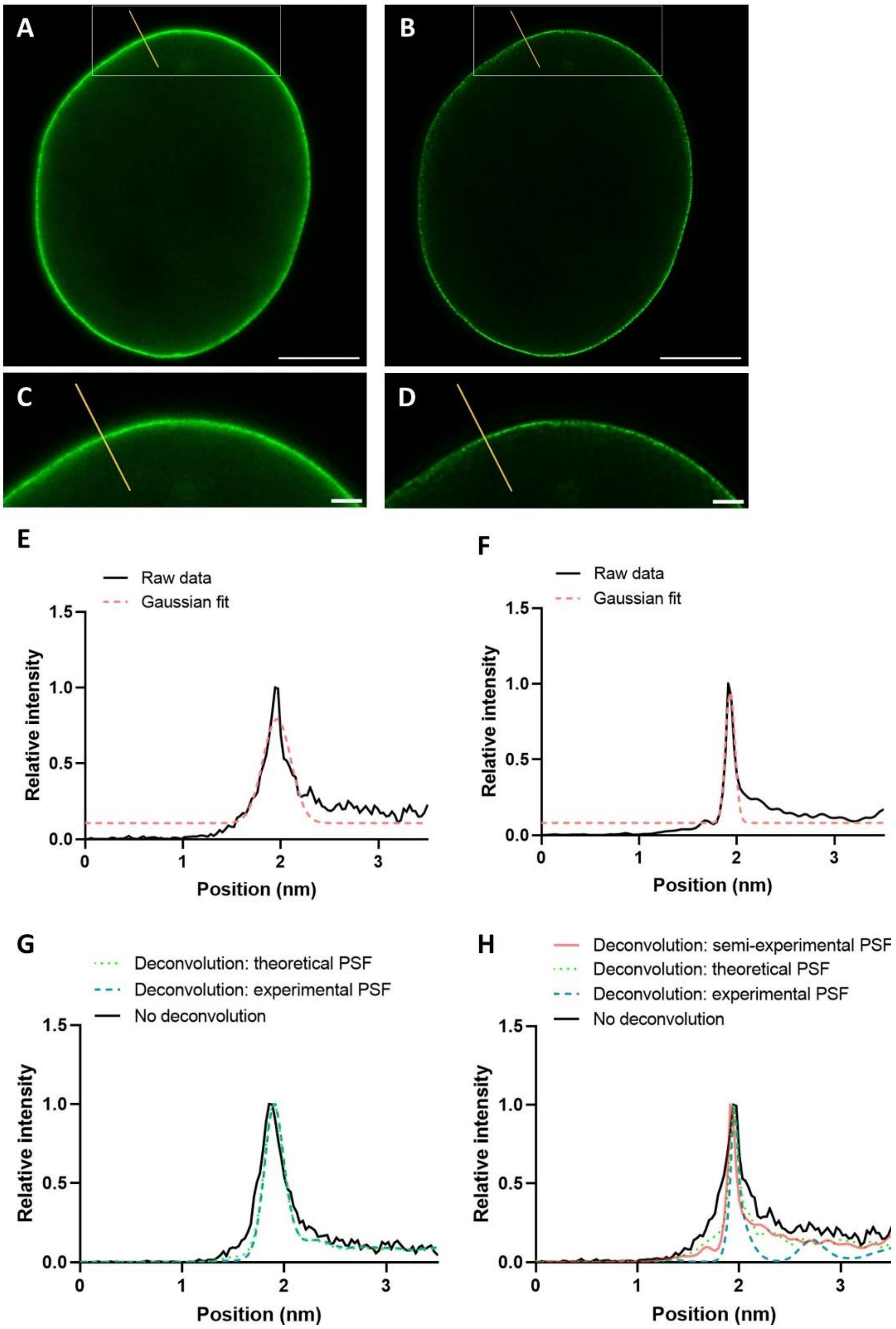


***Supplemental Figure 1:*** *Full Width Half Maximum (FHWM) determination in CSLM and STED microscopy images of NHDF stained for lamins A/C and B1. Only the lamins A/C staining is shown as example.* ***A-D****) STED microscopy image of lamins A/C, yellow line is used for the plot profiles displayed in (****E-G****);* ***A****) not deconvoluted,* ***B****) deconvoluted with semi-experimental PSF.* ***C-D****) Magnification of the ROI indicated in (****A****) and (****B****), respectively.* ***E****) Example of a Gaussian fit of non-deconvoluted STED microscopy image; Gaussian fit (pink dotted line) of the plot profile (black solid line) of the yellow line in (****A/C****). The height of the fitted curve is lower compared to the raw data and therefore the width at half of the peak will be broader than is truly the case in the raw data.* ***F****) Example of a Gaussian fit of deconvoluted STED microscopy image; Gaussian fit (pink dotted line) of the plot profile (black solid line) of the yellow line in (****B/D****).* ***G****) Plot profiles of CSLM, in images with no deconvolution (solid line) or deconvolution with the experimental PSF (blue dashed line), or theoretical (green dotted line).* ***H****) Plot profiles of STED microscopy, in images with no deconvolution (black solid line) or deconvolution with the experimental (blue dashed line), theoretical (green dotted line), or semi-experimental PSF (pink solid line). Scale bars A-B: 5 µm. Scale bars C-D: 1 µm*

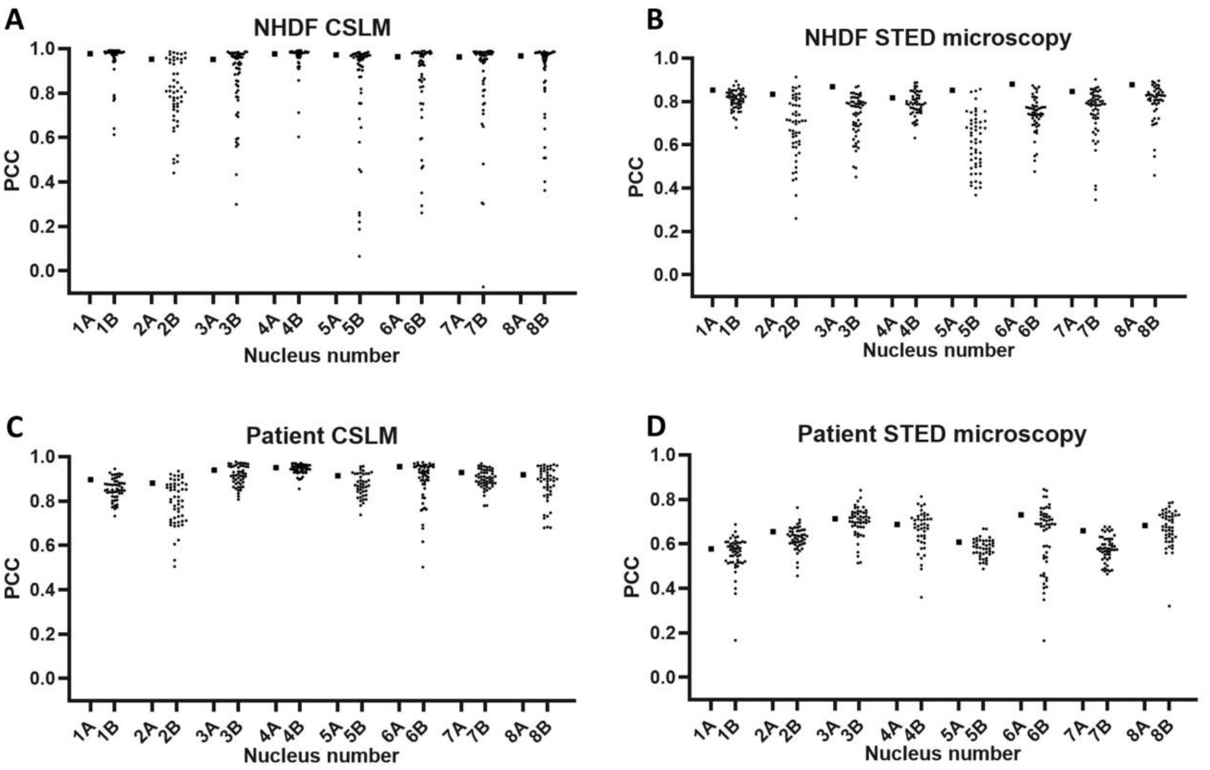


***Supplemental Figure 2:*** *Colocalization of lamins A/C and B1 expressed as Pearson's correlation coefficient (PCC) for the complete nucleus (one datapoint each for the 8 nuclei numbers indicated by A) or in the detailed approach (multiple datapoints for each of the 8 nuclei numbers indicated by B). For the detailed approach, microscopy images were divided into 100 equal rectangles and only rectangles with 100% overlap with the ROI of the nucleus were included in the colocalization analysis. (**A-B**) PCC of 8 CSLM (**A**) and 8 STED microscopy (**B**) images of NHDF. (**C-D**) PCC of 8 CSLM (**C**) and 8 STED microscopy (**D**) images of laminopathy patient dermal fibroblasts. The PCC was determined using Fiji.*
